# The Codon Statistics Database: a Database of Codon Usage Bias

**DOI:** 10.1101/2022.03.29.486291

**Authors:** Krishnamurthy Subramanian, Bryan Payne, Felix Feyertag, David Alvarez-Ponce

**Affiliations:** Biology Department, University of Nevada, Reno, Reno, NV, 89557; Department of Genetics, Rutgers, The State University of New Jersey, Piscataway, NJ, 08854

## Abstract

**Motivation:** Most amino acids can be encoded by a set of synonymous codons. Often, for any given amino acid, certain codons are significantly more used than others, a phenomenon known as codon usage bias. The genomes of different species differ in the frequencies at which they use each codon (e.g., a codon that is highly used in one species may be lowly used in another species). In addition, within any given genome, genes differ in their degree of codon bias, with highly expressed genes being more likely to use preferred codons. Knowing the codons that are preferred by a certain genome, and the amount of codon bias exhibited by each gene, has multiple applications (e.g., in heterologous expression, gene prediction, or phylogenetic inference).

**Results:** We have developed the Codon Statistics Database, an online database that contains codon usage statistics for all the species with reference or representative genomes in RefSeq. The user can search for any species and access two sets of tables. One set lists, for each codon, the frequency, the Relative Synonymous Codon Usage (RSCU), and whether the codon is preferred. Another set of tables lists, for each gene, its GC content, Effective Number of Codons (ENC), Codon Adaptation Index (CAI), and frequency of optimal codons (*F*_op_). Equivalent tables can be accessed for 1) all nuclear genes, 2) nuclear genes encoding ribosomal proteins, 3) mitochondrial genes and 4) chloroplastic genes (if available in the relevant assembly). The user can also search for any taxonomic group (e.g., “primates”) and obtain a table comparing all the species in the group.

**Availability:** The database is free to access without registration at http://codonstatsdb.unr.edu.

## INTRODUCTION

Most amino acids are encoded by multiple synonymous codons. Despite encoding for the same amino acid, some synonymous codons are used significantly more often than others, a phenomenon known as codon usage bias. Species significantly differ in their codon preferences – for instance, glutamic acid is preferentially encoded by GAG in human (Fig. 1), whereas the same amino acid is preferentially encoded by GAA in *Escherichia coli* (e.g. Sharp *et al*., 2010).

**Fig. 1.**
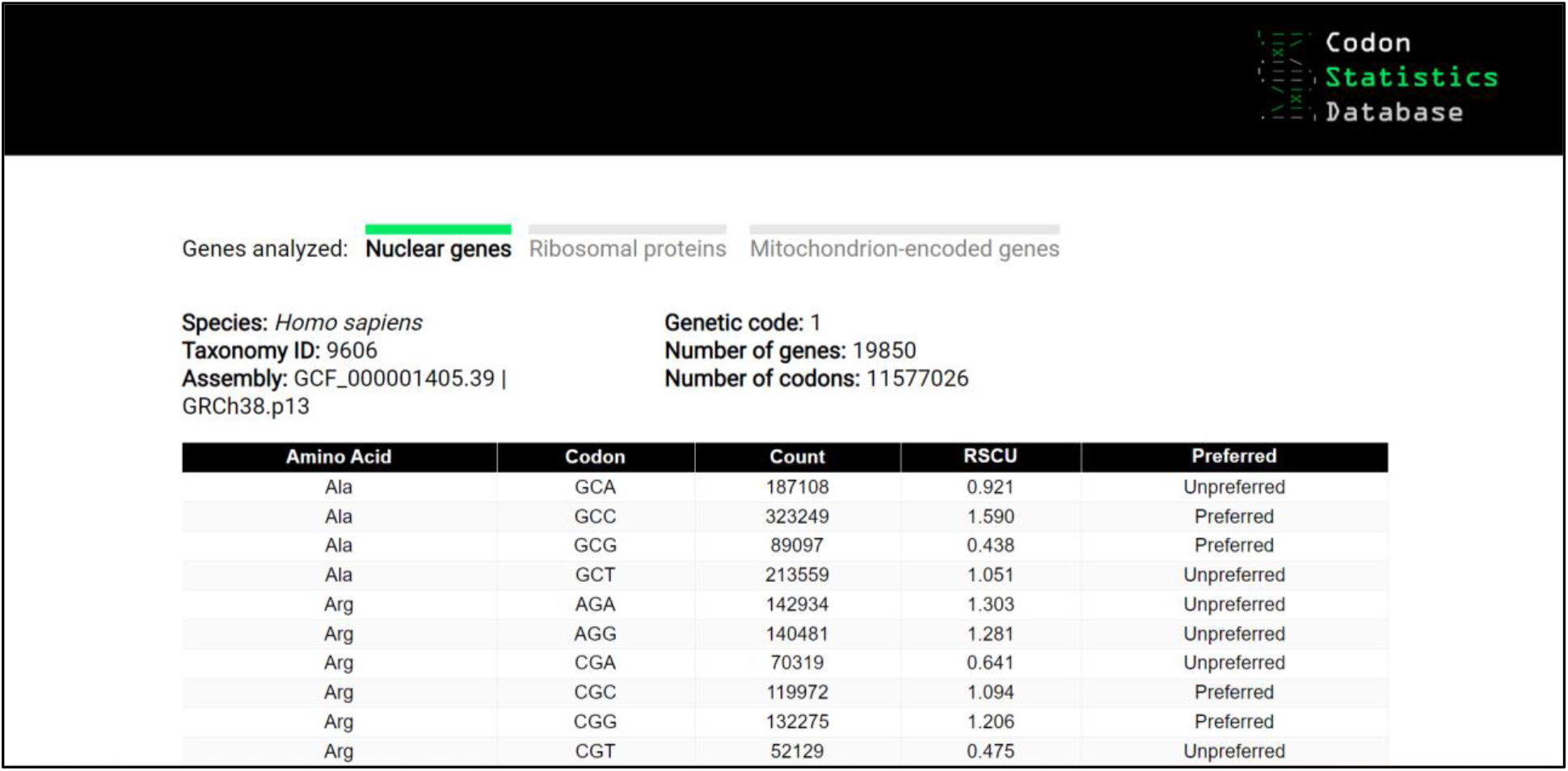
Species summary. Codon statistics corresponding to all human nuclear genes are shown.

In addition, genes within any given genome differ in their patterns of codon usage. In particular, gene expression levels significantly correlate with gene-specific metrics of codon usage such as the Effective Number of Codons (ENC; Wright, 1990), the Codon Adaptation Index (CAI; Sharp and Li, 1987) or the frequency of optimal codons (*F*_op_; Ikemura, 1985) (e.g., Gouy and Gautier, 1982).

Codon preferences can be affected by a number of factors, including the genome’s nucleotide composition (e.g., AT-rich genomes tend to use codons ending in A or T) and translational selection (codons that are translated by highly abundant tRNAs are translated faster and with fewer errors; e.g., Ikemura, 1985; Hershberg and Petrov, 2008).

Understanding codon preferences across the different species and genes is important not only to understanding genome evolution, but also for tasks such as heterologous expression, gene prediction or phylogenetic inference (e.g., Gustafsson *et al*., 2004; Christianson, 2005). In addition, the patterns of codon usage of viruses tend to be similar to those of their host species (e.g., Shackelton *et al*., 2006).

Despite the relevance of maintaining species- and gene-specific codon usage information, existing databases are deprecated, limited in phylogenetic scope, and/or do not provide gene-specific metrics (Nakamura *et al*. 2000; Hilterbrand *et al*., 2012; Athey *et al*., 2017).

## IMPLEMENTATION

For each of the species with reference or representative genomes in the RefSeq database (release 207), we chose one full assembly (in order of preference, the one labeled as “reference”, the one with the highest assembly level, or the most recent one) and retrieved the corresponding CDSs file. Using that file as input, a number of tables were pre-computed using an R pipeline. Only one CDS per gene was used (if multiple were available, the longest one was chosen). The web interface was created using PERL CGI.

For each species, we computed the total frequency of each codon, and used the information to compute the Relative Synonymous Codon Usage (RSCU) of each codon. For each gene, we computed the GC content for the entire CDS (GC), the GC content at third codon positions (GC3), the ENC and the RSCU for each codon.

For species with over 1000 genes, we also compared genes inferred to be highly expressed (bottom 10% ENC values) with genes inferred to be lowly expressed (top 10% ENC values). Codons with significantly higher RSCU values in the highly expressed gene set (according to a Mann-Whitney *U* test) were considered preferred/optimal. We then computed the *F*_op_ for each gene. The highly expressed gene set was also used as reference to compute the CAI of each gene.

## THE DATABASE

We have created the Codon Statistics Database, an online database that contains codon usage information for all species with reference or representative genomes in RefSeq (over 15,000). The user can search for any species or taxonomic group by taxonomic ID (e.g. “9606”), scientific name (e.g. “*Homo sapiens*”) or common name (e.g. “human”), and select an option from a drop-down menu.

If a species is selected, the user is directed to a table that lists, for each codon, the encoded amino acid, the total count in the genome, the RSCU, and whether the codon is preferred or unpreferred (Fig. 1). The user can access equivalent tables for 1) all nuclear genes (default option), 2) nuclear genes encoding ribosomal proteins (this subset is included since such proteins are often highly expressed and thus subjected to strong codon bias), 3) mitochondrion-encoded genes and 4) chloroplast-encoded genes (if such gene sets are available in the relevant genome assembly). For viruses, only one option including all genes is available. Additionally, for each gene set, the user can download a tab-delimited file (.tsv) listing the following statistics for each gene: GC, GC3, ENC, CAI, and *F*_op_.

If a taxonomic group with multiple species is selected (e.g. “7215”, “*Drosophila*” or “fruit flies”), the user is presented with a table comparing all the species in the group (Fig. 2). The user has the option to visualize either codon counts or RSCU values. Preferred codons in each species are marked with asterisks.

**Fig. 2.**
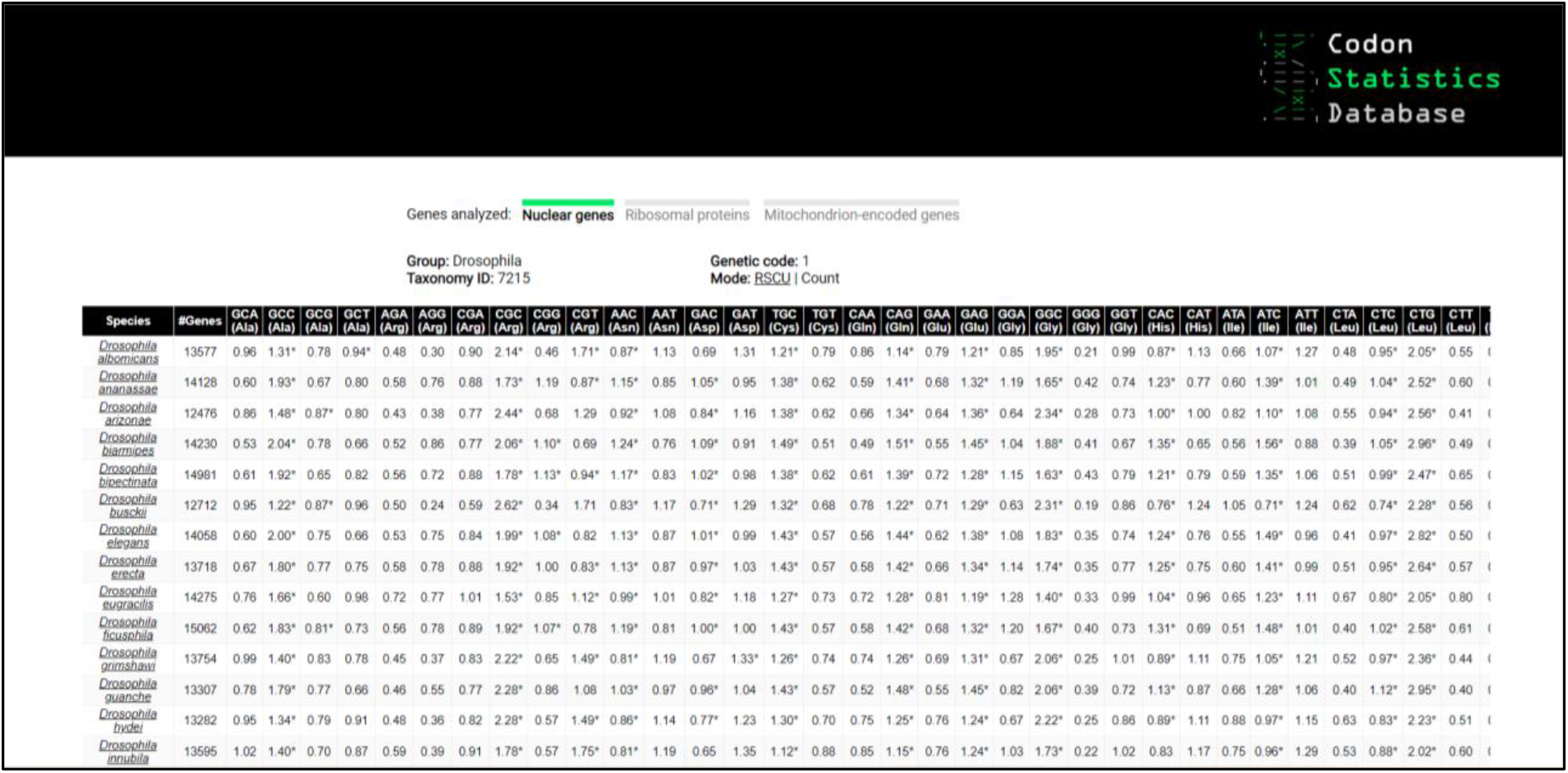
Taxonomic group summary. Codon preferences for species in the genus *Drosophila* are shown.

## ACHNOWLEDGEMENTS

We are grateful to Alejandra Nores for help with web design, and to the University of Nevada, Reno Information Technology Operations for computational resources.

